## Supplemental tabels and figures for "30-Minute Highly Multiplexed VaxArray Immunoassay for Pneumococcal Vaccine Antigen Characterization"

**Supplemental Table 1**: Reactivity of conjugates using 23-Mix Primary Detection Label. Signal to blank ratios (S/Bl) generated on the VaxArray 23-valent Pneumococcal Assay for monovalent Pfizer- and EuBiologics (EuB)-provided conjugates at 2 µg/mL using the 23-Mix Primary Detection Label. Columns represent each serotype-specific capture antibody, and rows represent each conjugate tested. Text in bold green shows S/Bl>3 generated for the matched conjugate and capture antibody. Text in bold red shows S/Bl<3 generated for the matched conjugate and capture antibody indicating low reactivity. Text in bold black shows S/Bl>3 generated on off-target capture antibody indicating cross-reactivity.

| | | **Serotype-Specific Microarry Capture Antibody** | | | | | | | | | | | | | | | | | | | | | | |
|---|---|---|---|---|---|---|---|---|---|---|---|---|---|---|---|---|---|---|---|---|---|---|---|---|
| | | **S1** | **S2** | **S3** | **S4** | **S5** | **S6A** | **S6B** | **S7F** | **S8** | **S9N** | **S9V** | **S10A** | **S12F** | **S14** | **S15B** | **S17F** | **S18C** | **S19A** | **S19F** | **S20** | **S22F** | **S23F** | **S33F** |
| Pfizer | S1 | **5.8** | 1.0 | 1.0 | 1.0 | 0.9 | 1.0 | 0.9 | 0.9 | 0.9 | 1.0 | 1.0 | 0.9 | 1.0 | 1.0 | 1.0 | 1.0 | 0.9 | 0.9 | 1.1 | 1.1 | 1.1 | 1.1 | 1.2 |
| EuB | | **12.6** | 1.1 | 0.9 | 1.0 | 0.9 | 0.9 | 0.7 | 0.6 | 0.5 | 0.8 | 0.9 | 0.8 | 0.6 | 0.6 | 0.7 | 0.7 | 0.7 | 0.6 | 0.8 | 1.0 | 1.1 | 0.9 | 1.0 |
| Pfizer | S3 | 0.9 | 1.1 | **6.8** | 1.0 | 0.9 | 1.0 | 0.9 | 1.0 | 0.9 | 0.9 | 0.9 | 0.9 | 1.0 | 1.0 | 1.1 | 1.0 | 1.0 | 1.0 | 1.0 | 1.1 | 1.1 | 1.0 | 1.3 |
| EuB | | 1.1 | 1.1 | **7.7** | 1.2 | 1.1 | 1.0 | 1.1 | 1.1 | 1.1 | 1.0 | 1.0 | 1.1 | 1.1 | 1.2 | 1.2 | 1.3 | 1.2 | 1.1 | 1.0 | 1.1 | 1.7 | 1.1 | 1.3 |
| Pfizer | S4 | 1.0 | 1.3 | 1.0 | **9.3** | 1.1 | 1.1 | 1.0 | 1.0 | 1.0 | 1.0 | 1.0 | 1.0 | 1.1 | 1.0 | 1.1 | 1.0 | 1.1 | 1.1 | 1.1 | 1.2 | 1.1 | 1.1 | 1.3 |
| EuB | | 1.2 | 1.3 | 1.3 | **9.9** | 1.2 | 1.2 | 1.2 | 1.4 | 1.1 | 1.1 | 1.1 | 1.2 | 1.2 | 1.1 | 1.4 | 1.4 | 1.3 | 1.1 | 1.1 | 1.3 | 1.1 | 1.2 | 1.4 |
| Pfizer | S5 | 1.0 | 1.1 | 1.0 | 0.9 | **11.8** | 1.0 | 0.9 | 0.9 | 1.2 | 1.0 | 1.0 | 1.0 | 1.0 | 1.0 | 1.0 | 1.0 | 0.9 | 0.9 | 1.0 | 1.1 | 1.1 | 1.1 | 1.1 |
| EuB | | 1.2 | 1.3 | 1.2 | 1.3 | **12.9** | 1.3 | 1.2 | 1.2 | 1.5 | 1.1 | 1.1 | 1.1 | 1.2 | 1.2 | 1.3 | 1.3 | 1.3 | 1.2 | 1.2 | 1.1 | 1.2 | 1.1 | 1.3 |
| Pfizer | S6A | 1.1 | 1.0 | 1.0 | 0.9 | 0.9 | **8.3** | 1.0 | 0.9 | 0.9 | 0.9 | 1.0 | 1.1 | 1.0 | 1.0 | 1.0 | 0.9 | 0.9 | 0.9 | 1.1 | 1.2 | 1.1 | 1.1 | 1.1 |
| EuB | | 1.1 | 1.2 | 1.2 | 1.2 | 1.1 | **7.7** | 1.1 | 1.1 | 1.0 | 1.0 | 1.0 | 1.1 | 1.0 | 1.0 | 1.2 | 1.1 | 1.1 | 1.0 | 1.0 | 1.0 | 1.0 | 1.0 | 1.1 |
| Pfizer | S6B | 1.1 | 1.2 | 1.1 | 1.1 | 1.1 | 1.2 | **8.9** | 1.1 | 1.1 | 1.1 | 1.0 | 1.0 | 1.0 | 1.1 | 1.2 | 1.1 | 1.2 | 1.1 | 1.0 | 1.1 | 1.0 | 1.0 | 1.2 |
| EuB | | 1.1 | 1.2 | 1.1 | 1.1 | 1.0 | 1.2 | **8.7** | 1.1 | 1.0 | 1.1 | 1.0 | 1.0 | 1.0 | 1.0 | 1.1 | 1.1 | 1.2 | 1.0 | 1.0 | 1.2 | 1.0 | 1.1 | 1.2 |
| Pfizer | S7F | 1.0 | 1.0 | 0.9 | 1.1 | 1.0 | 1.1 | 1.0 | **9.9** | 1.0 | 1.0 | 1.0 | 1.1 | 1.0 | 1.0 | 1.0 | 1.0 | 1.0 | 1.1 | 1.0 | 1.2 | 1.0 | 1.0 | 1.2 |
| EuB | | 1.1 | 1.1 | 1.2 | 1.2 | 1.1 | 1.2 | 1.1 | **13.6** | 1.1 | 1.0 | 1.0 | 1.1 | 1.0 | 1.0 | 1.1 | 1.1 | 1.3 | 1.0 | 1.0 | 1.0 | 1.0 | 0.9 | 1.0 |
| Pfizer | S9V | 1.0 | 1.1 | 0.9 | 1.0 | 1.0 | 1.1 | 1.0 | 1.0 | 1.0 | 1.0 | **14.3** | 1.2 | 1.0 | 1.0 | 1.0 | 1.0 | 1.0 | 1.1 | 1.0 | 1.0 | 1.0 | 0.9 | 1.2 |
| EuB | | 1.3 | 1.8 | 1.7 | 1.9 | 1.2 | 1.4 | 1.3 | 1.1 | 1.2 | 1.3 | **10.3** | 1.3 | 1.4 | 1.2 | 1.7 | 1.8 | 1.6 | 1.1 | 1.1 | 1.9 | 1.3 | 2.0 | 2.0 |
| Pfizer | S14 | 1.6 | 1.1 | 0.9 | 0.9 | 1.0 | 1.1 | 1.0 | 1.0 | 1.0 | 1.0 | 1.0 | 1.1 | 1.1 | **11.8** | 1.0 | 1.0 | 1.1 | 1.1 | 1.0 | 1.1 | 1.0 | 0.8 | 1.3 |
| EuB | | 2.0 | 2.2 | 1.7 | 2.2 | 1.3 | 1.6 | 1.5 | 1.2 | 1.3 | 1.4 | 1.3 | 1.5 | 1.4 | **14.9** | 2.1 | 2.0 | 1.9 | 1.3 | 1.1 | 2.1 | 1.4 | 2.1 | 2.1 |
| Pfizer | S18C | 1.0 | 1.0 | 0.9 | 1.0 | 1.0 | 1.0 | 1.0 | 1.0 | 0.9 | 1.0 | 1.0 | 1.0 | 1.0 | 1.0 | 1.0 | 1.0 | **1.1** | 1.0 | 1.0 | 1.0 | 1.0 | 0.9 | 1.1 |
| EuB | | 1.4 | 2.0 | 1.6 | 1.9 | 1.3 | 1.5 | 1.4 | 1.1 | 1.1 | 1.4 | 1.2 | 1.3 | 1.4 | 1.2 | 1.9 | 2.0 | **13.7** | 1.2 | 1.2 | 1.8 | 1.3 | 2.0 | 2.1 |
| Pfizer | S19A | 1.0 | 0.9 | 0.8 | 0.8 | 1.0 | 0.9 | 1.0 | 1.0 | 0.9 | 1.0 | 1.0 | 1.0 | 1.0 | 1.0 | 0.9 | 0.9 | 1.0 | **9.3** | 1.0 | 0.9 | 1.0 | **5.9** | 1.0 |
| EuB | | 1.4 | 2.0 | 1.7 | 2.0 | 1.3 | 1.5 | 1.4 | 1.1 | 1.3 | 1.4 | 1.3 | 1.3 | 1.4 | 1.2 | 1.7 | 1.8 | 1.7 | **16.1** | 1.1 | 1.8 | 1.3 | 1.8 | 1.8 |
| Pfizer | S19F | 1.0 | 0.9 | 1.0 | 1.0 | 0.9 | 0.9 | 1.0 | 1.0 | 0.9 | 1.0 | 0.9 | 1.1 | 1.1 | 0.9 | 1.2 | 1.1 | 1.1 | 1.0 | **12.7** | 1.3 | 1.0 | 1.1 | 1.3 |
| EuB | | 1.5 | 2.2 | 1.9 | 2.1 | 1.3 | 1.6 | 1.4 | 1.2 | 1.2 | 1.5 | 1.3 | 1.4 | 1.4 | 1.2 | 1.7 | 1.7 | 1.9 | 1.2 | **16.1** | 1.9 | 1.4 | 1.9 | 1.9 |
| EuB | S22F | 0.9 | 0.9 | 0.9 | 0.9 | 0.9 | 0.9 | 0.9 | 0.9 | 0.9 | 0.9 | 0.9 | 0.9 | 0.9 | 0.9 | 0.8 | 0.9 | 0.9 | 0.9 | 0.9 | 0.9 | **10.5** | 0.9 | 0.9 |
| Pfizer | S23F | 0.9 | 0.8 | 0.9 | 0.9 | 0.9 | 1.0 | 1.0 | 0.9 | 0.9 | 1.0 | 0.9 | 0.9 | 1.0 | 0.9 | 1.1 | 1.2 | 1.1 | 1.0 | 0.9 | 1.1 | 1.0 | **5.2** | 1.2 |
| EuB | | 0.9 | 1.0 | 1.0 | 0.9 | 0.9 | 0.9 | 0.9 | 0.9 | 0.9 | 0.8 | 0.9 | 0.9 | 0.9 | 1.0 | 0.9 | 0.9 | 0.9 | 0.9 | 0.9 | 0.9 | 0.9 | **3.3** | 1.0 |

**Supplemental Table 2**: Reactivity of conjugates using anti-CRM197 Primary Detection Label. Signal to blank ratios (S/Bl) generated on the VaxArray 23-valent Pneumococcal Assay for monovalent Pfizer- and EuBiologics (EuB)-provided conjugates at 2 µg/mL labeled with anti-CRM197 Primary Detection Label. Columns represent each serotype-specific capture antibody, and rows represent each conjugate tested. Text in bold green shows S/Bl>3 generated for the matched conjugate and capture antibody. Text in bold black shows S/Bl>3 generated on off-target capture antibody indicating cross-reactivity.

| | | **Serotype-Specific Microarry Capture Antibody** | | | | | | | | | | | | | | | | | | | | | | |
|---|---|---|---|---|---|---|---|---|---|---|---|---|---|---|---|---|---|---|---|---|---|---|---|---|
| | | **S1** | **S2** | **S3** | **S4** | **S5** | **S6A** | **S6B** | **S7F** | **S8** | **S9N** | **S9V** | **S10A** | **S12F** | **S14** | **S15B** | **S17F** | **S18C** | **S19A** | **S19F** | **S20** | **S22F** | **S23F** | **S33F** |
| Pfizer | S1 | **12.0** | 1.0 | 1.0 | 1.0 | 1.0 | 1.0 | 1.0 | 1.0 | 1.0 | 1.0 | 1.0 | 1.0 | 1.0 | 1.0 | 1.0 | 1.0 | 1.0 | 1.0 | 1.0 | 1.0 | 1.0 | 1.0 | 1.0 |
| EuB | | **5.8** | 1.0 | 1.0 | 1.0 | 1.0 | 1.0 | 1.0 | 1.0 | 1.0 | 1.0 | 1.0 | 1.1 | 1.0 | 1.1 | 1.0 | 1.0 | 1.0 | 1.0 | 1.0 | 1.0 | 1.1 | 1.0 | 1.0 |
| Pfizer | S3 | 1.1 | 1.0 | **13.2** | 1.0 | 1.0 | 1.0 | 1.0 | 1.0 | 1.0 | 1.1 | 1.0 | 1.0 | 1.0 | 1.0 | 1.0 | 1.0 | 1.1 | 1.0 | 1.0 | 1.0 | 1.0 | 1.0 | 1.0 |
| EuB | | 1.1 | 1.1 | **11.9** | 1.1 | 1.1 | 1.1 | 1.1 | 1.1 | 1.1 | 1.1 | 1.1 | 1.1 | 1.1 | 0.9 | 1.1 | 1.1 | 1.1 | 1.1 | 1.1 | 1.0 | 1.7 | 1.1 | 1.0 |
| Pfizer | S4 | 1.1 | 1.0 | 1.1 | **18.1** | 1.1 | 1.1 | 1.0 | 1.0 | 1.0 | 1.1 | 1.1 | 1.1 | 1.1 | 1.1 | 1.1 | 1.1 | 1.1 | 1.0 | 1.1 | 1.1 | 1.0 | 1.1 | 1.1 |
| EuB | | 1.1 | 1.0 | 1.0 | **12.1** | 1.1 | 1.1 | 1.0 | 1.1 | 1.0 | 1.1 | 1.0 | 1.1 | 1.1 | 1.2 | 1.0 | 1.1 | 1.0 | 1.0 | 1.1 | 1.0 | 1.1 | 1.1 | 1.0 |
| Pfizer | S5 | 1.0 | 1.0 | 1.1 | 1.1 | **13.4** | 1.1 | 1.0 | 1.0 | 1.1 | 1.1 | 1.0 | 1.0 | 1.0 | 0.9 | 1.0 | 1.0 | 1.0 | 1.0 | 1.0 | 1.0 | 1.0 | 1.0 | 1.0 |
| EuB | | 1.1 | 1.0 | 1.0 | 1.0 | **3.6** | 1.0 | 1.0 | 1.0 | 1.1 | 1.0 | 1.0 | 1.0 | 1.1 | 1.1 | 1.0 | 1.0 | 1.0 | 1.0 | 1.0 | 1.0 | 1.0 | 1.0 | 1.0 |
| Pfizer | S6A | 1.0 | 1.0 | 1.0 | 1.0 | 1.0 | **12.4** | 1.0 | 1.0 | 1.0 | 1.1 | 1.0 | 1.0 | 1.0 | 1.0 | 1.0 | 1.0 | 1.0 | 1.0 | 1.1 | 1.0 | 1.0 | 1.0 | 1.0 |
| EuB | | 1.0 | 1.0 | 1.0 | 1.0 | 1.1 | **11.5** | 1.1 | 1.0 | 1.0 | 1.0 | 1.1 | 1.0 | 1.0 | 1.1 | 1.0 | 1.0 | 1.0 | 1.0 | 1.1 | 1.0 | 1.0 | 1.0 | 1.0 |
| Pfizer | S6B | 1.0 | 1.0 | 1.0 | 1.0 | 1.0 | 0.9 | **12.5** | 0.9 | 1.0 | 1.0 | 1.0 | 1.0 | 1.0 | 1.1 | 0.9 | 1.0 | 1.0 | 1.0 | 1.0 | 1.0 | 1.0 | 1.0 | 1.0 |
| EuB | | 1.0 | 1.0 | 1.0 | 1.0 | 1.0 | 1.1 | **6.3** | 1.0 | 1.0 | 1.0 | 1.0 | 1.0 | 1.0 | 1.0 | 1.0 | 1.0 | 1.0 | 1.0 | 0.9 | 1.0 | 1.0 | 1.0 | 1.0 |
| Pfizer | S7F | 1.0 | 1.0 | 1.0 | 1.0 | 1.0 | 1.0 | 1.0 | **10.1** | 1.0 | 1.0 | 1.0 | 0.9 | 1.0 | 0.8 | 1.0 | 1.0 | 1.0 | 1.0 | 1.0 | 1.0 | 0.9 | 1.0 | 1.0 |
| EuB | | 1.0 | 1.0 | 1.0 | 1.0 | 1.0 | 1.0 | 1.0 | **8.7** | 1.0 | 1.0 | 1.0 | 1.0 | 1.0 | 1.0 | 1.0 | 1.0 | 1.1 | 1.0 | 1.0 | 1.0 | 1.0 | 1.0 | 1.0 |
| Pfizer | S9V | 0.9 | 0.9 | 0.9 | 1.0 | 1.0 | 0.9 | 0.9 | 0.9 | 0.9 | 0.9 | **17.9** | 1.0 | 1.0 | 1.0 | 0.9 | 0.9 | 1.0 | 0.9 | 0.9 | 1.0 | 1.0 | 0.9 | 1.0 |
| EuB | | 1.0 | 1.0 | 1.0 | 1.0 | 1.0 | 1.0 | 1.0 | 1.0 | 1.0 | 0.9 | **15.5** | 1.0 | 1.0 | 1.1 | 1.0 | 1.0 | 1.0 | 1.0 | 1.0 | 1.0 | 1.0 | 1.0 | 1.0 |
| Pfizer | S14 | 1.0 | 1.0 | 1.0 | 1.0 | 1.0 | 1.0 | 1.0 | 1.0 | 1.0 | 1.0 | 1.0 | 1.0 | 1.0 | **7.0** | 1.0 | 1.0 | 1.0 | 1.0 | 0.9 | 1.0 | 1.0 | 0.9 | 0.9 |
| EuB | | 1.1 | 1.0 | 0.9 | 1.0 | 1.0 | 1.0 | 1.0 | 1.0 | 1.0 | 0.9 | 1.0 | 1.0 | 0.9 | **5.7** | 1.0 | 1.0 | 1.0 | 0.9 | 0.9 | 1.0 | 1.0 | 1.0 | 0.9 |
| Pfizer | S18C | 1.0 | 1.0 | 1.0 | 1.0 | 1.0 | 1.0 | 1.0 | 1.0 | 1.0 | 0.9 | 1.0 | 1.0 | 1.0 | 0.9 | 1.1 | 1.0 | **13.6** | 1.0 | 1.0 | 1.0 | 0.9 | 1.0 | 1.0 |
| EuB | | 1.0 | 1.0 | 1.0 | 1.0 | 1.1 | 1.0 | 1.0 | 1.1 | 1.0 | 1.0 | 1.0 | 1.0 | 1.0 | 1.1 | 1.0 | 1.1 | **8.3** | 1.0 | 1.0 | 1.0 | 1.0 | 1.1 | 1.0 |
| Pfizer | S19A | 1.0 | 0.9 | 1.0 | 1.0 | 1.0 | 1.0 | 0.9 | 0.9 | 1.0 | 1.0 | 1.0 | 0.9 | 0.9 | 1.0 | 0.9 | 0.9 | 0.9 | **12.9** | 0.9 | 1.0 | 1.0 | **13.4** | 1.0 |
| EuB | | 0.9 | 1.0 | 0.9 | 1.0 | 1.0 | 1.0 | 0.9 | 1.0 | 1.0 | 0.9 | 0.9 | 0.9 | 1.0 | 1.0 | 1.0 | 1.0 | 0.9 | **7.0** | 1.0 | 0.9 | 0.9 | 1.0 | 0.9 |
| Pfizer | S19F | 1.0 | 1.0 | 1.0 | 1.0 | 1.0 | 1.0 | 1.0 | 1.0 | 1.0 | 1.0 | 1.0 | 1.0 | 1.0 | 1.0 | 1.0 | 1.0 | 1.0 | 1.0 | **9.9** | 0.9 | 1.0 | 0.9 | 1.0 |
| EuB | | 1.0 | 1.0 | 0.9 | 0.9 | 1.0 | 0.9 | 1.0 | 1.0 | 0.9 | 1.0 | 0.9 | 1.1 | 1.0 | 1.0 | 0.9 | 1.0 | 1.0 | 0.9 | **5.5** | 1.0 | 1.0 | 1.0 | 0.9 |
| EuB | S22F | 1.0 | 1.0 | 1.0 | 0.9 | 0.9 | 0.9 | 0.9 | 0.9 | 0.9 | 0.9 | 0.9 | 1.0 | 0.9 | 0.8 | 0.9 | 0.9 | 0.9 | 0.9 | 0.9 | 0.9 | **6.0** | 0.9 | 0.9 |
| Pfizer | S23F | 1.0 | 1.0 | 1.0 | 1.0 | 1.0 | 1.0 | 1.0 | 1.0 | 1.0 | 1.0 | 1.0 | 0.9 | 1.0 | 0.9 | 1.0 | 1.0 | 1.0 | 1.0 | 0.9 | 1.0 | 1.0 | **16.0** | 1.1 |
| EuB | | 0.9 | 0.9 | 0.9 | 0.9 | 0.9 | 0.9 | 1.0 | 0.9 | 0.9 | 0.9 | 0.9 | 0.9 | 0.9 | 0.9 | 0.9 | 0.9 | 0.9 | 0.9 | 0.9 | 0.9 | 0.9 | **8.8** | 0.9 |

**Supplemental Table 3**: Analytical sensitivity and working range for 13-valent Pfizer conjugates and 14-valent EuBiologics (EuB) conjugates with both 23-Mix and anti-CRM197 Primary Detection Labels. "NA": not applicable due to low signal on S18C capture.

| Serotype | Supplier | **Approx. LLOQ** (ng/mL) | | **Approx ULOQ** (µg/mL) | | **Working Range** (x-fold) | |
|---|---|---|---|---|---|---|---|
| | | **23-Mix Label** | **anti-CRM197 Label** | **23-Mix Label** | **anti-CRM197 Label** | **23-Mix Label** | **anti-CRM197 Label** |
| **1** | Pfizer | 2.4 | 8.6 | 0.4 | 0.8 | 181 | 95 |
| | EuB | 3.7 | 13.5 | 0.8 | 0.8 | 226 | 59 |
| **3** | Pfizer | 4.3 | 6.7 | 0.6 | 0.7 | 144 | 97 |
| | EuB | 28.4 | 14.8 | 1.8 | 1.6 | 64 | 109 |
| **4** | Pfizer | 2.5 | 4.2 | 0.5 | 0.5 | 204 | 123 |
| | EuB | 8.5 | 17.7 | 0.8 | 1.1 | 100 | 62 |
| **5** | Pfizer | 8.6 | 7.8 | 0.8 | 0.8 | 94 | 106 |
| | EuB | 43.0 | 98.9 | 1.7 | 1.7 | 39 | 17 |
| **6A** | Pfizer | 47.3 | 27.4 | 2.4 | 1.9 | 51 | 71 |
| | EuB | 45.5 | 11.7 | 2.0 | 1.4 | 45 | 118 |
| **6B** | Pfizer | 76.3 | 26.6 | 6.6 | 3.5 | 86 | 130 |
| | EuB | 78.0 | 12.8 | 2.5 | 1.9 | 31 | 148 |
| **7F** | Pfizer | 14.4 | 38.2 | 3.0 | 3.3 | 207 | 87 |
| | EuB | 20.2 | 39.7 | 1.8 | 2.2 | 90 | 55 |
| **9V** | Pfizer | 29.8 | 8.9 | 1.7 | 1.2 | 57 | 129 |
| | EuB | 190.7 | 14.3 | 2.5 | 1.6 | 13 | 110 |
| **14** | Pfizer | 2.5 | 43.8 | 1.0 | 1.7 | 385 | 40 |
| | EuB | 8.2 | 41.4 | 1.1 | 2.0 | 129 | 49 |
| **18C** | Pfizer | NA | 8.9 | NA | 0.6 | NA | 69 |
| | EuB | 18.0 | 23.8 | 2.1 | 1.5 | 115 | 63 |
| **19A** | Pfizer | 23.9 | 7.8 | 1.6 | 1.1 | 66 | 146 |
| | EuB | 49.0 | 14.5 | 1.9 | 1.5 | 39 | 104 |
| **19F** | Pfizer | 13.6 | 27.3 | 1.4 | 1.1 | 106 | 41 |
| | EuB | 15.5 | 30.7 | 1.1 | 1.3 | 72 | 44 |
| **22F** | EuB | 14.1 | 29.0 | 1.8 | 1.3 | 127 | 43 |
| **23F** | Pfizer | 26.2 | 6.1 | 1.5 | 1.0 | 56 | 167 |
| | EuB | 128.1 | 9.1 | 2.6 | 1.2 | 21 | 127 |

**Supplemental Table 4:** Accuracy and precision for EuBiologics 15-valent drug product with and without sodium citrate desorption using 23-Mix Primary Detection Label. Text in red indicates values not within 80-120% recovery and < 20% RSD.

| Serotype | Expected Conc. (µg/mL) | No Desorption | | Desorbed | |
|---|---|---|---|---|---|
| | | Accuracy (% Recovery) | Precision (%RSD) | Accuracy (% Recovery) | Precision (%RSD) |
| **S1** | 0.18 | 116% | 8% | 118%* | 14%* |
| **S3** | 0.16 | 104% | 10% | 110% | 13% |
| **S4** | 0.18 | **79%** | **24%** | 100%* | 12%* |
| **S5** | 0.32 | 109% | 19% | 83% | 19% |
| **S6A** | 0.28 | 94% | 8% | 97% | 13% |
| **S6B** | 0.67 | 96% | 9% | 105% | 11% |
| **S7F** | 0.20 | 92% | 11% | 104% | 15% |
| **S9V** | 0.20 | 115% | 12% | 102% | 19% |
| **S14** | 0.13 | 94% | 17% | 88% | 7% |
| **S18C** | 0.12 | **26%** | 6% | 84% | 11% |
| **S19A** | 0.25 | 93% | 5% | 101% | 11% |
| **S19F** | 0.13 | 113% | **27%** | 91% | 6% |
| **S22F** | 0.24 | 97% | 8% | 111% | 11% |
| **S23F** | 0.27 | 111% | **65%** | 106% | 9% |
| **Overall** | | 96±23% | 16±16% | 98±10% | 12±4% |

*: Data is from 400ms exposure, Std 1 signal saturated at 700ms.

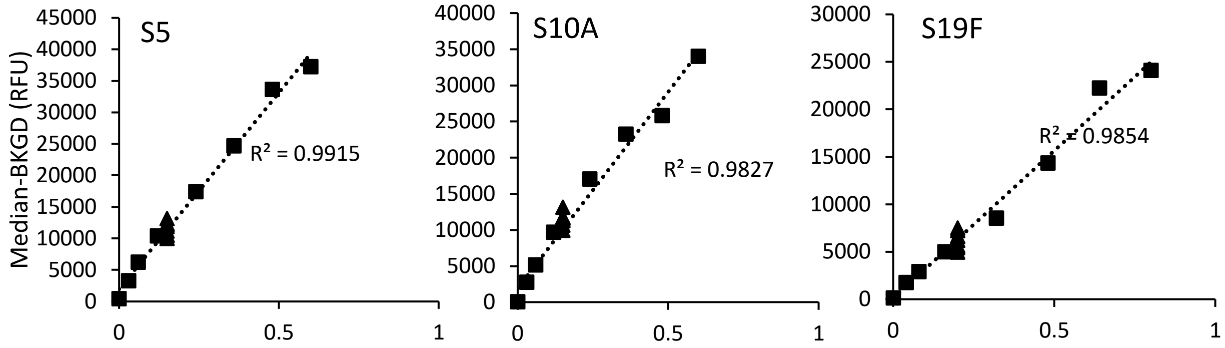


**Supplemental Figure 1.** Signal response (median signal with background signal subtracted) of samples measured (triangles) and matched standard (squares) for Serotype 5, 10A and 19F in 23-valent Pfizer native sample. Linear fits are dotted lines with the associated correlation efficient indicated.

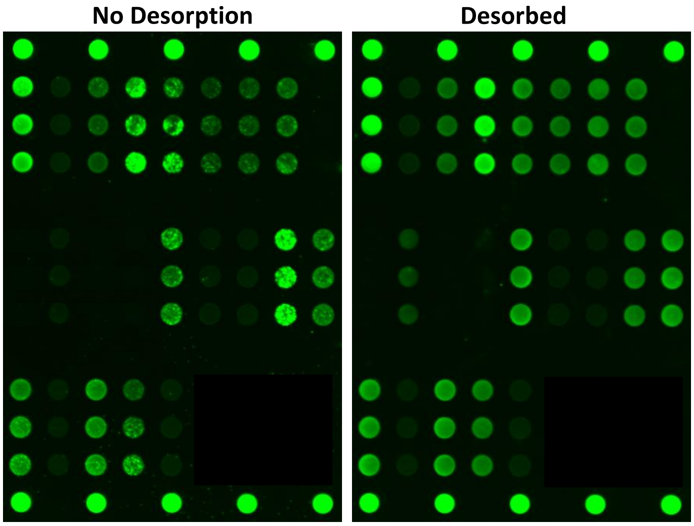


**Supplemental Figure 2.** Representative fluorescence microarray images of samples under quantification before desorption (left) and post-desorption (right) showing improved microarray spot morphologies.
